# Investigation of neuromodulatory effect of anodal cerebellar transcranial direct current stimulation on the primary motor cortex using functional near-infrared spectroscopy

**DOI:** 10.1101/2022.09.30.510216

**Authors:** Zeshan Shoaib, Won Kee Chang, Jongseung Lee, Stephanie Hyeyoung Lee, V Zephaniah Phillips, Seung Hyun Lee, Nam-Jong Paik, Han-Jeong Hwang, Won-Seok Kim

## Abstract

**Background:** Cerebellar brain inhibition (CBI), a neural connection between the cerebellum and primary motor cortex (M1), has been researched as a target pathway for neuromodulation to improve clinical outcomes in various neurological diseases. However, conflicting results of anodal cerebellar transcranial direct current stimulation (acb-tDCS) on M1 excitability indicate that additional investigation is required to examine its precise effect.

**Objective/Hypothesis:** This study aimed to gather evidence of the neuromodulatory effect of acb-tDCS on the M1 using functional near-infrared spectroscopy (fNIRS).

**Methods:** Sixteen healthy participants were included in this cross-over study. Participants received real and sham acb-tDCS in a random order, with a minimum one-week washout period between them. The anode and cathode were placed on the right cerebellum and the right buccinator muscle, respectively. Stimulation lasted 20 min at an intensity of 2 mA, and fNIRS data were recorded for 42 min (including a 4 min baseline before stimulation and an 18 min post-stimulation duration) using eight channels attached bilaterally on the M1.

**Results:** acb-tDCS induced a significant decrease in oxyhemoglobin (HbO) concentration (inhibitory effect) in the left (contralateral) M1, whereas it induced a significant increase in HbO concentration (excitatory effect) in the right (ipsilateral) M1 compared to sham tDCS during (*p* < 0.05) and after stimulation (*p* < 0.01) in a group level analysis. At the individual level, variations in the response to acb-tDCS were observed.

**Conclusion:** Our findings demonstrate the neuromodulatory effects of acb-tDCS on the bilateral M1 in terms of neuronal hemodynamics.

## Introduction

When a negative event is likely to happen in daily life, the human brain generates a negative predictive error signal [1, 2], which triggers extinction neural mechanisms. This adaptive inhibitory learning process gradually suppresses and eventually eliminates maladaptive behaviors, which may lead to abnormal activation of mossy fiber inputs to the cerebellum or extensive destabilization of cerebellar circuits, resulting in general inconsistencies in motor performance [3]. These findings suggest that the cerebellum, through neural connections with the motor cortex, plays an important role in motor learning [4]. Purkinje cells are activated in the cerebellar cortex and these activities are transmitted to the dentate nucleus, which inhibits the M1 [5]. This procedure is referred to as cerebellar brain inhibition (CBI). The concept of CBI is fairly developed from previous research; however, more research is required to determine the precise effect of modulating cerebellar excitability on the M1.

In recent years, transcranial direct current stimulation (tDCS) has emerged as a non-invasive brain stimulation technique that can modulate neural activity and thereby improve brain functions in healthy as well as patients [6, 7]. It induces excitability changes in the human brain that are bidirectional (excitatory or inhibitory) based on input polarity (anodal or cathodal) [8, 9]. Although tDCS was initially applied to investigate cerebral cortical function [10], due to the cortical and subcortical connections of the cerebellum, many researchers have also started to use cerebellar stimulation in a variety of preclinical and clinical studies [11, 12]. The field of cerebellar-tDCS (cb-tDCS) has recently received much consideration in clinical research, especially because it may provide novel insights into cerebellar neural plasticity [13]. It has also been discovered that cb-tDCS can modulate excitability in the deep cerebellar output nuclei by polarizing the Purkinje cells [14, 15].

Many researchers have investigated the effect of cb-tDCS on CBI but have reported inconsistent results; some studies have found that cb-tDCS decreases the effect of CBI [16, 17], while others have found the opposite [15]. Because the cerebellar cortical Purkinje cells have connections with the M1, it is possible that changes in the excitability of Purkinje cells may also induce changes in M1 excitability. Therefore, previous CBI studies investigated the effects of cb-tDCS on M1 excitability, but the results were also inconsistent. In 2009, Galea et al. demonstrated that anodal cb-tDCS (acb-tDCS) has an excitatory effect on the M1 [15], whereas some studies found no effect after acb-tDCS [18]. Interestingly, Ates et al. showed in 2018 that acb-tDCS has both excitatory and inhibitory effects on M1 excitability [19]. Moreover, Liebrand et al. recently showed an increase in ipsilateral M1 activation following acb-tDCS [20]. These conflicting results may be because the cb-tDCS parameters used in previous studies have been quite heterogenous across the studies and there is a lack of data regarding how the cerebellar stimulation precisely affects the variety of neuronal populations, such as Purkinje cells and granule cells [21].

Among various neuroimaging modalities for studying the metabolic state of the brain, such as functional magnetic resonance imaging (fMRI), single-photon emission computerized tomography, and positron emission tomography (PET), the emerging neuroimaging modality of functional near-infrared spectroscopy (fNIRS) may help to address the challenge of measuring neuronal activity in real-time with reasonable spatial and better temporal resolution [22, 23]. fNIRS has additional advantages over fMRI and PET, such as portability, low cost, and better safety [24]. fNIRS provides metabolic information non-invasively and with adequate sensitivity to detect even small changes in the cerebral hemodynamic response. It allows estimation of metabolic-based specificity [25], and robustness to various artifacts [26]. Furthermore, fNIRS can be used simultaneously with tDCS to measure brain activity even during the stimulation period without significant electro-optic interference.

The current study aimed to investigate M1 cortical activity through HbO concentration changes caused by acb-tDCS using fNIRS. We used a sham-controlled, cross-over study design for anodal and sham cb-tDCS during the resting state. We used eight (located on the contralateral and ipsilateral M1) of the 56 fNIRS channels to investigate cortical excitability in terms of HbO concentration changes, where the anode and cathode were positioned on the right cerebellum and the right buccinator muscle, respectively. We compared the excitability changes of the contralateral and ipsilateral M1 under real and sham conditions. The hemodynamic changes on both sides of the M1 were monitored not only before and after the stimulation, but also during the stimulation delivery itself. Since tDCS-mediated modulation of dentate nucleus activity could change the M1 cortical excitability, we hypothesized that acb-tDCS would decrease the hemodynamic response of the contralateral M1 (inhibitory effect) in comparison to sham tDCS. To better understand the precise effect of acb-tDCS on the M1, we also investigated the inter-subject variability results along with the group level analysis.

## Material and methods

### Participants

Inclusion criteria for this study were healthy individuals aged 19 or older, who were right-handed as assessed by the Edinburgh handedness inventory [27], were able to understand the study protocol, and who gave written consent. The exclusion criteria were the following: any history of disease involving the central nervous system such as stroke, traumatic brain injury or Parkinson’s disease; and any history of the following conditions: scalp lesions interfering with tDCS application, pregnancy, cardiac pacemakers, cochlear implants, or metals in the head (e.g., clip, coil). Initially, 20 subjects (12 men, 8 women; mean age 26 ± 3 years) were enrolled. Due to excessive noisy channels, we excluded 4 subjects’ data from analysis, leaving 16 subjects (9 men, 7 women; mean age 26 ± 3 years) for data analysis. All participants were informed about the study procedures and possible risks and provided written consent prior to the experiment. This study was carried out according to the Declaration of Helsinki and Good Clinical Practice Guidelines, and the protocol was approved by the Seoul National University Bundang Hospital Institutional Review Board (IRB No. B-1908-558-004).

### fNIRS instrumentation and configuration

The functional neuromodulations were investigated using a continuous wave fNIRS system, NIRScout (NIRx Medical technologies, LLC., MN, USA). The system operates in the near-infrared spectrum at two different wavelengths (760 nm and 850 nm) with a sampling rate of 3.91 Hz. Sixteen sources and 20 detectors were employed, which created 56 channels, and the approximate distance between the sources and detectors was maintained at 3 cm using the NIRScaps (NIRx Medical technologies, LLC., MN, USA) [28]. The optode placement robustness among all subjects was confirmed by conducting proper adjustments and measurements. Because the study aimed to monitor the neuromodulation due to cb-tDCS in the M1, we only considered NIRS channels around C3, Cz, and C4 locations based on the EEG 10-20 international system for data analysis. Therefore, among the 56 channels, eight channels were used for further data analysis (Figure 1; four on the contralateral hemisphere and four on the ipsilateral hemisphere).

**Figure 1.**
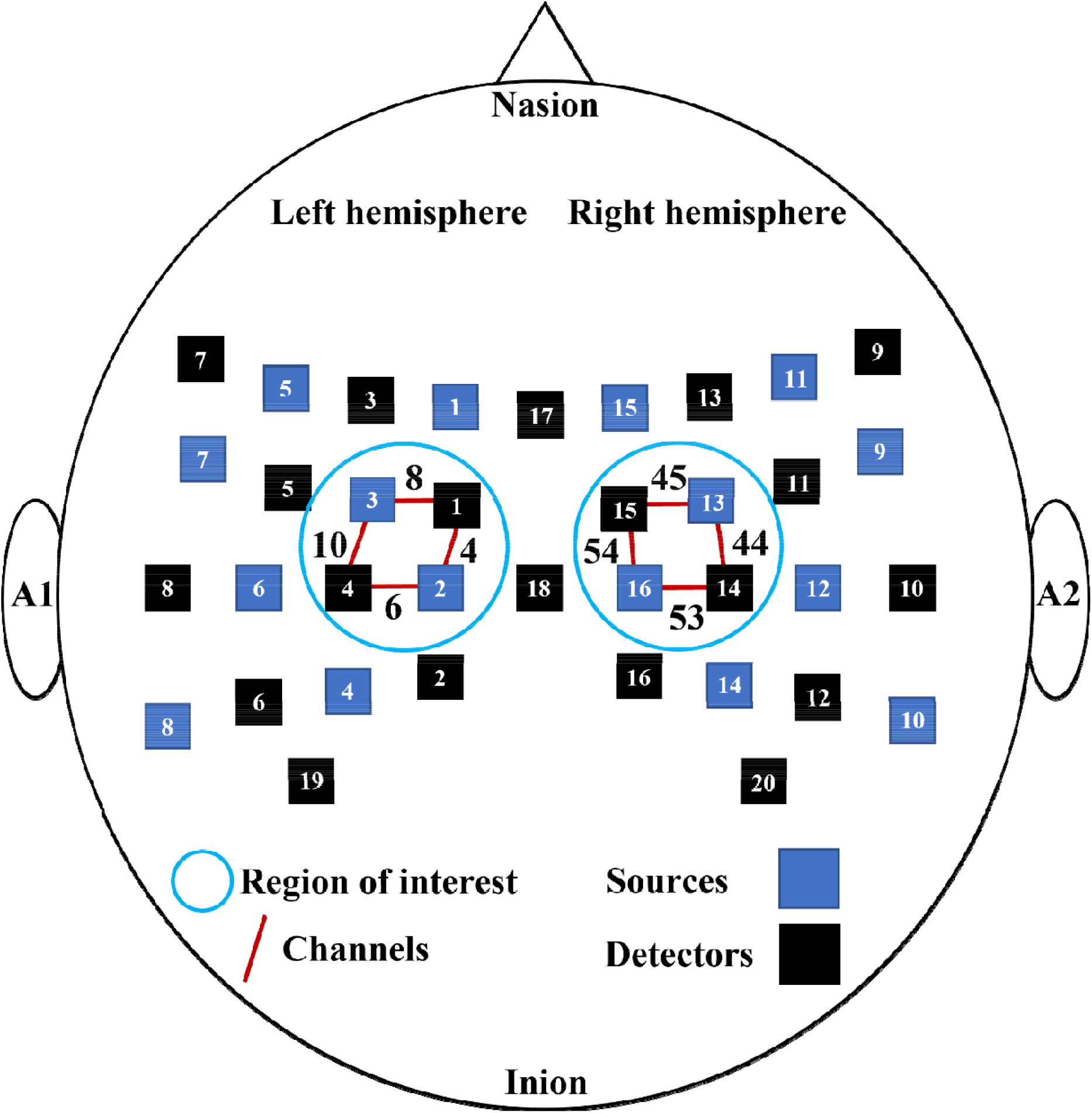
The optode arrangement of the 16 sources (marked in blue boxes) and 20 detectors (marked in black boxes) on the subject’s head used for data acquisition yielded 56 active NIRS channels. Blue circles represent the two ROIs in the left and right hemispheres, and targeted channels are labeled with black numbers on the red lines. Detector 18 is placed on the Cz position of the international 10-20 system.

### Cerebellar-tDCS and experimental design

A single channel tDCS device was used for the experiment (MINDD STIM, Ybrain, Korea). Two electrodes of a 60 mm diameter were used with saline-soaked sponge pads to increase conduction. We positioned the anodal electrode on the right cerebellar cortex (3-4 cm lateral to the inion) and the cathode electrode on the ipsilateral buccinator muscle [15, 29]. The electrodes were firmly attached using a head band. We used the auto alarm system which was activated when high impedance occurred to maintain secure electrode attachment. The same procedure was utilized for both real and sham tDCS.

The experimental paradigm comprised three phases: a 4-min pre-stimulation period, a 20-min stimulation period, and an additional 18-min post-stimulation period. Real-stimulation experiments provided 20 min of anodal stimulation, which was ramped up to 2.0 mA over 30 s, maintained at 2.0 mA for 19 min, then ramped down to 0 mA over 30 s. For sham tDCS experiments, the current was ramped up to 2 mA over 30 s, ramped down over the next 30 s, and then stopped. All subjects underwent real and sham stimulation experiments at least seven days apart in a random order. The participants and researchers were blinded to the stimulation condition (real or sham). tDCS was provided at rest in both experiments, and participants were not required to be involved in any motor task. Each participant was seated in a comfortable chair with their eyes closed but was instructed not to fall asleep throughout the experiment.

### Preprocessing and data analysis

MATLAB R2021b (MathWorks, Inc., Natick, MA, USA) software was used to analyze the NIRS data. Optical intensities were converted to oxygenated and deoxygenated hemoglobin (HbO/HbR) concentration changes by the Modified Beer-Lambert law [22, 30] and saved in the MATLAB workspace for further processing. An age- and wavelength-dependent differential pathlength factor was calculated for each subject [31]. In this study, the HbR time course showed insignificant changes in comparison to the HbO time course throughout the experiment [32, 33]; therefore, we only focused on HbO concentration changes. fNIRS signals were filtered using a Butterworth low-pass filter (4^th^ order) with a cut-off frequency of 0.1 Hz [32]. Subsequently, the filtered-NIRS data were used to compute the mean HbO concentration changes in the contralateral and ipsilateral hemispheres for each subject. In the end, subject-wise, channel-wise, and hemisphere-wise grand-averaged HbO concentration changes were computed.

### Statistical analysis

Two tests were used to verify the normality and homogeneity of the variances of distribution: the Kolmogorov–Smirnov test and Levene’s test, respectively. We used the parametric paired t-test and the non-parametric Wilcoxon signed rank test for the data that did and did not meet normality, respectively. These statistical tests were conducted for subject-wise, channel-wise, and hemisphere-wise comparisons of real and sham stimulations. All statistical tests used a 95% confidence interval, implying that a *p*-value of less than 0.05 was significant. We quantified the inhibitory and excitatory effects by displaying the statistical significance between the HbO levels of the real and sham stimulation.

## Results

### Spatiotemporal representation of hemodynamic responses

To analyze the effects of both real and sham cb-tDCS on HbO, the time course was divided into three different time windows: before (baseline period), during (stimulation period), and after (post-stimulation period) stimulation. Figures 2(a) and (b) depict the channel-wise grand-averaged time course of HbO on the contralateral and ipsilateral hemispheres, respectively. The HbO concentration decreased after real cb-tDCS stimulation initiation as compared to the sham stimulation of the contralateral hemisphere (inhibitory effect), whereas the opposite result was observed in the ipsilateral hemisphere (excitatory effect). This phenomenon was also clearly represented in the hemisphere-wise grand-average time course of HbO, as shown in Figure 2(c). Figure 3 shows the spatiotemporal representation of hemodynamic responses, which also clearly presents the mentioned inhibitory and excitatory effects of acb-tDCS on the contralateral and ipsilateral hemispheres, respectively. Overall, real acb-tDCS (lower row in each box) led to a significant increase in HbO concentration in the ipsilateral hemisphere and decrease in HbO concentration in the contralateral hemisphere. However, sham tDCS (upper row in each box) showed relatively little change in HbO concentration over time.

**Figure 2.**
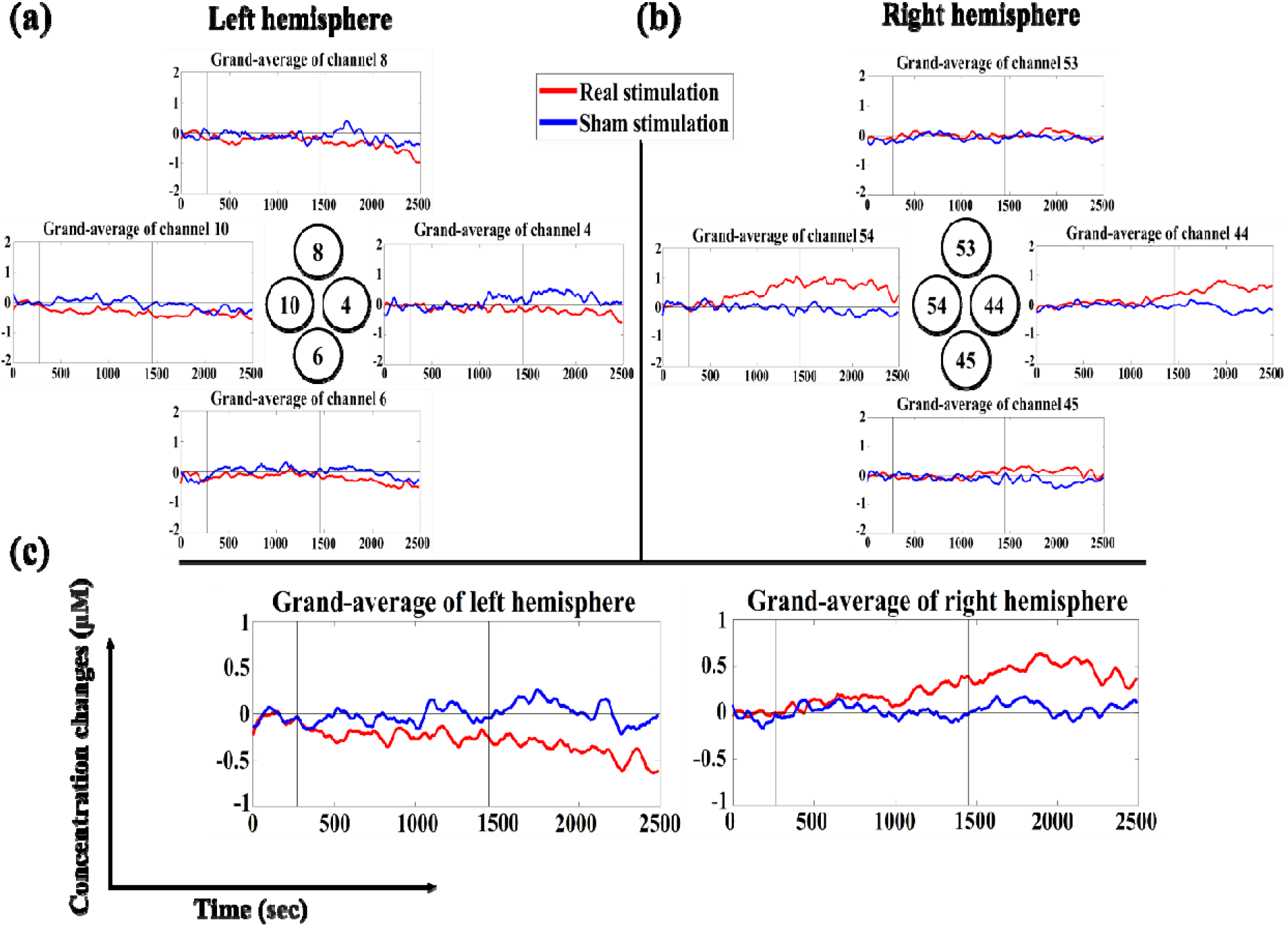
Grand-averaged HbO concentration changes of **(a)** channels 4, 6, 8, and 10 on the left hemisphere and **(b)** channels 44, 45, 53, and 54 on the right hemisphere over all subjects, and **(c)** those of the left and right hemispheres. The real and sham tDCS are denoted by red and blue lines, respectively. The x-axis represents the time in seconds, and the y-axis represents the HbO concentration changes in micro mole units. The first vertical line represents the end of the baseline duration, and the second vertical line represents the end of the stimulation duration.

**Figure 3.**
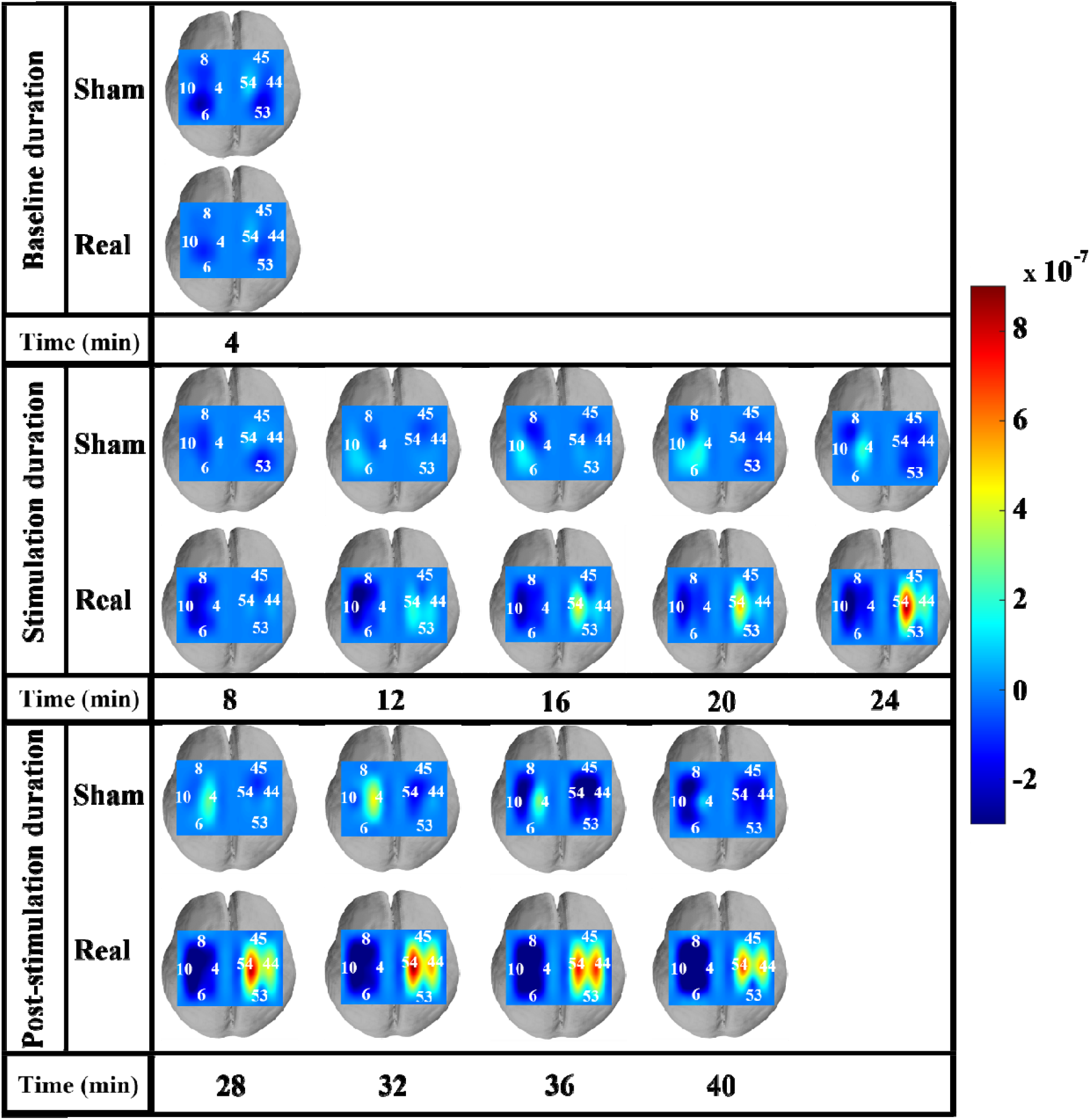
Spatiotemporal pattern maps of sham (upper in each box) and real (lower) cb-tDCS. Each box shows topographical brain maps for all channels on the left and right hemispheres with an interval of 4 min. Each topographical map was obtained by averaging HbO concentrations for every 4 min.

### Quantification of hemodynamic responses

Figure 4 presents quantified hemodynamic responses for channel- and hemisphere-wise grand-average hemodynamic responses. No statistical difference was observed between real and sham tDCS in the baseline period (see “B” in Figure 4) for all results. However, the inhibitory and excitatory effects were statistically significant in the stimulation and post-stimulation periods for most channels of the contralateral (see “S” and “P” in Figure 4(a)) and ipsilateral (see “S” and “P” in Figure 4(b)) hemispheres, respectively (*p* < 0.05). Some channels did not show statistically significant differences between real and sham tDCS, but did show the inhibitory and excitatory effects for the corresponding hemisphere with marginally statistical results, i.e., channel 8 (*p* = 0.123 for the stimulation period (“S”)), channel 45 (*p* = 0.071 for the stimulation period (“S”)), and channel 53 (*p* = 0.093 for the stimulation period (“S”)). The statistical trend of the channel-wise grand-average hemodynamic response is clearly represented in the hemisphere-wise grand-average ones (Figure 4(c)); significant inhabitation and excitation were observed in the stimulation (“S”) (*p* < 0.05) and post-stimulation (“P”) (*p* < 0.01) periods for the contralateral and ipsilateral hemispheres, respectively.

**Figure 4.**
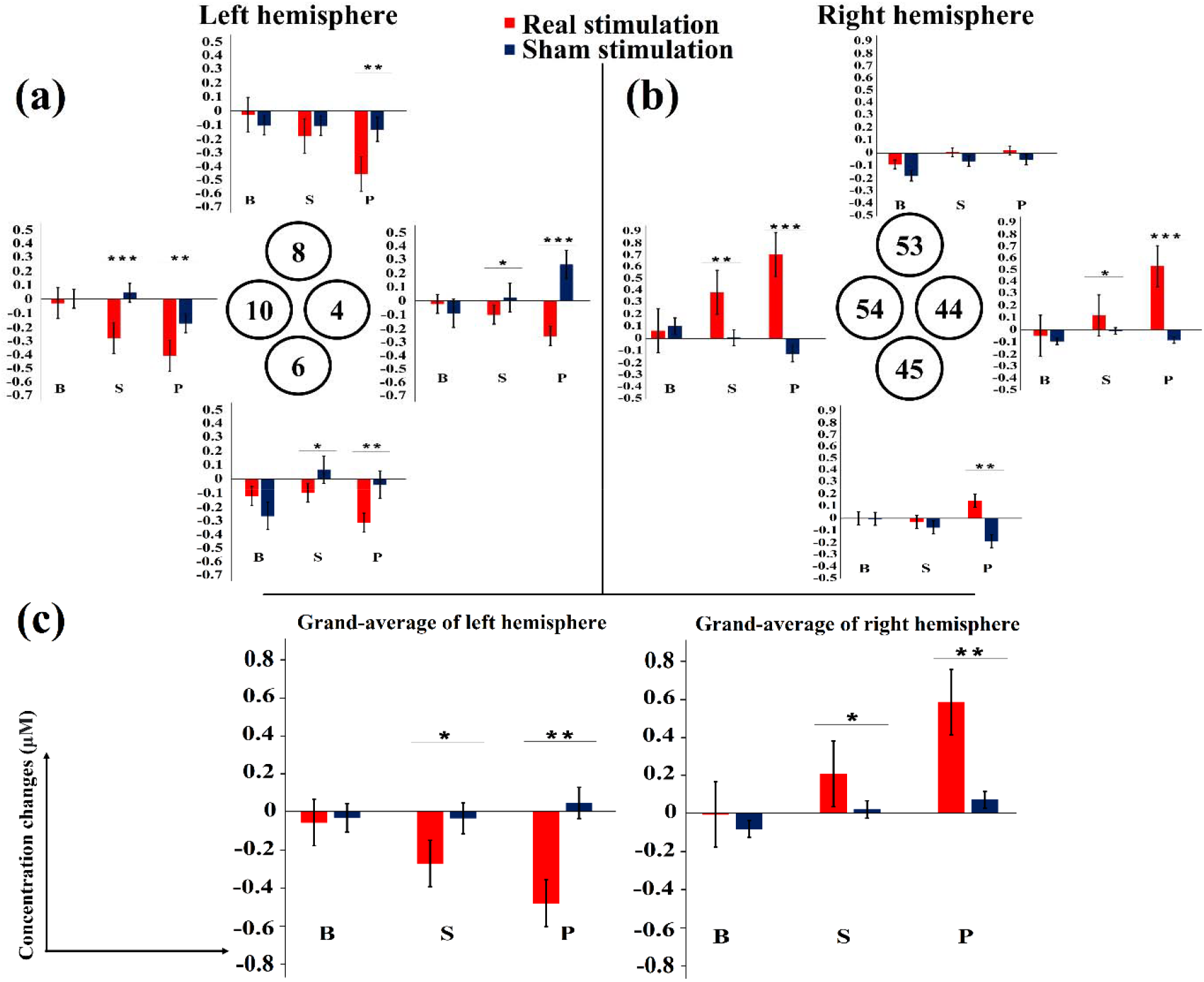
Channel-wise and hemisphere-wise mean changes in HbO concentration during B (baseline), S (stimulation), and P (post-stimulation) periods of the real (red bars) and sham (blue bars) cb-tDCS for each channel on the **(a)** left (channels 4, 6, 8, and 10) and **(b)** right (channels 44, 45, 53, and 54) hemispheres. **(c)** Grand-averaged HbO concentration changes over the left and right hemispheres. The y-axis represents the HbO concentration changes in micro mole units. The error bars represent standard errors of the mean values; significance is marked as follows: \**p*□<□0.05; \*\**p*□<□0.01; \*\*\**p*□<□0.001.

### Individual variability in response to acb-tDCS

Figure 5 shows the mean HbO concentration values of each subject in the three different phases of the real and sham tDCS on the contralateral M1. Four subjects (25%) failed to show an inhibitory response to acb-tDCS (subjects 4, 7, 8, and 10) and two of them (subjects 4 and 7) showed excitatory responses. Three of the subjects (subjects 2, 12, and 16) who had an inhibitory response showed an abnormal increase during sham tDCS, while the other nine subjects (56%) showed inhibitory behavior with a normal sham response. Similarly, Figure 6 presents each subject’s mean HbO concentration values in three different phases of the real and sham tDCS on the ipsilateral M1. Five subjects (31%) failed to exhibit an excitatory response to acb-tDCS (subjects 1, 2, 11, 12, and 14). Two of these subjects (12 and 14) showed increasing trends for real acb-tDCS, but they had a lower HbO concentration than sham tDCS due to an abnormal increase during sham tDCS. One subject (subject 1) showed an inhibitory response to acb-tDCS on the ipsilateral M1; the other two subjects (subjects 2 and 11) showed little change during acb-tDCS.

**Figure 5.**
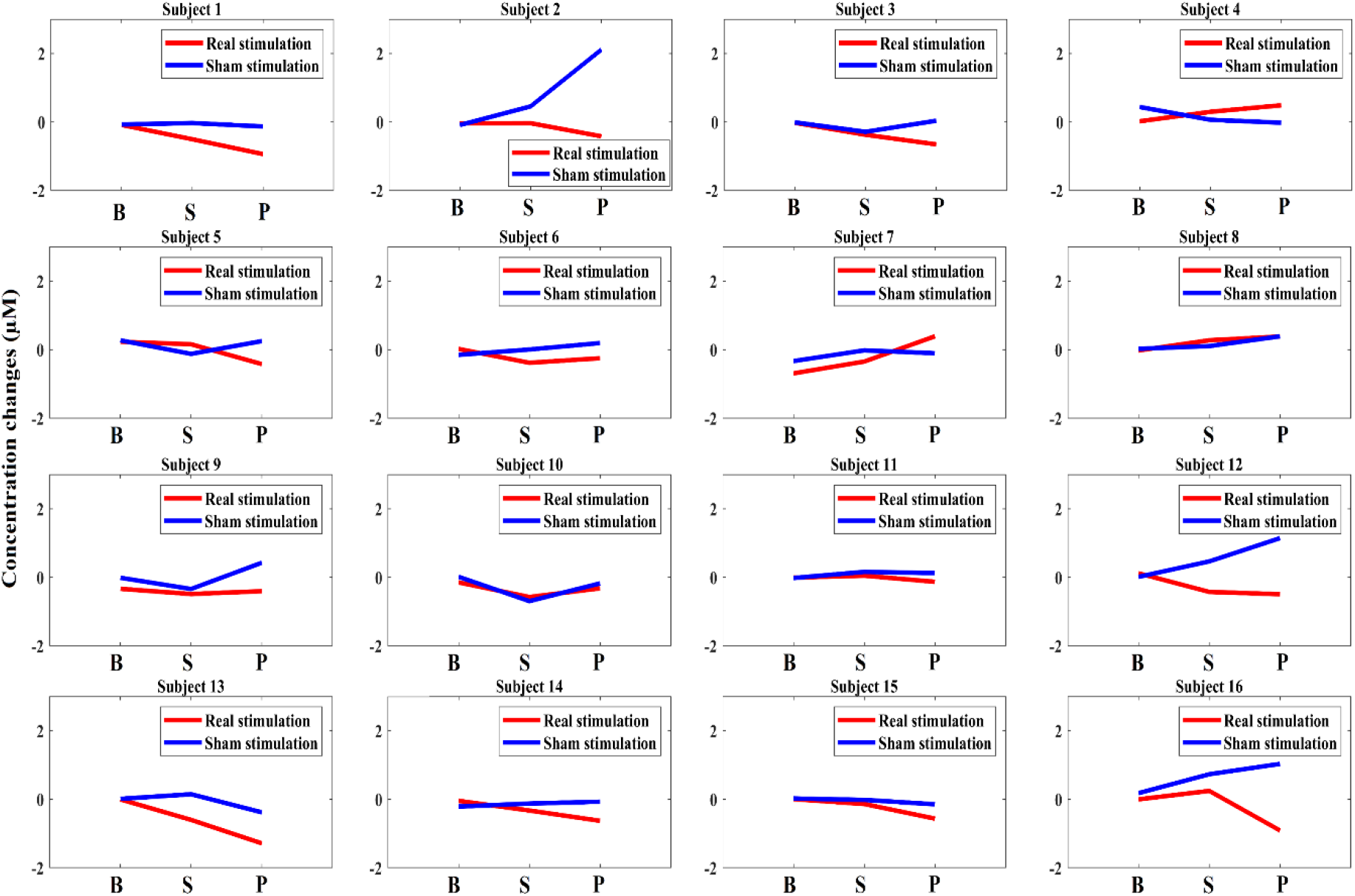
Mean concentration changes of HbO in three different phases for each subject’s real (red line) and sham (blue line) cb-tDCS on the left M1. On the x-axis, B, S, and P represent the average HbO concentration in baseline, stimulation, and post-stimulation periods, respectively.

**Figure 6.**
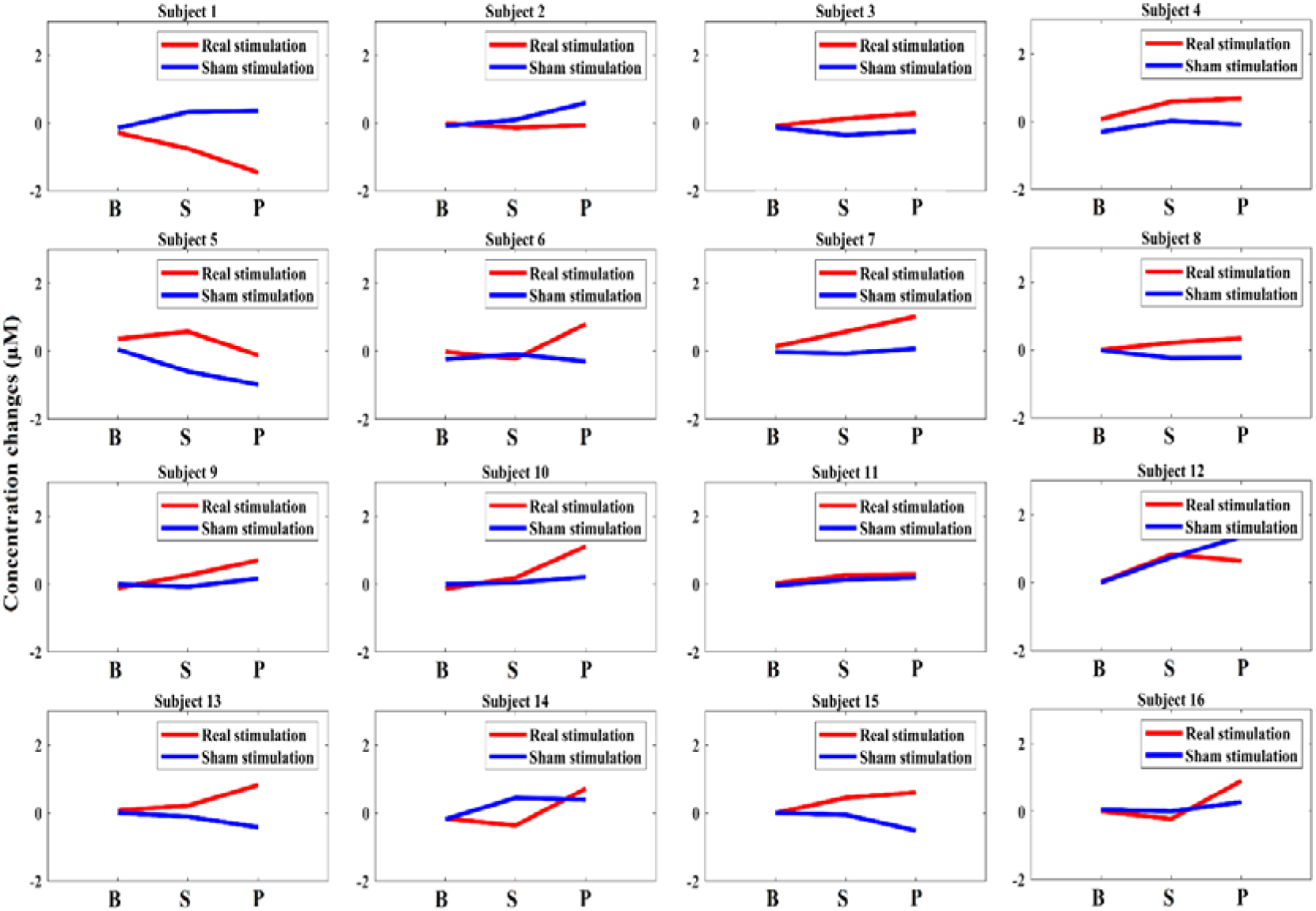
Mean concentration changes of HbO in three different phases for each subject’s real (red line) and sham (blue line) cb-tDCS on the right M1. On the x-axis, B, S, and P represent the average HbO concentration in baseline, stimulation, and post-stimulation periods, respectively.

## Discussion

The main findings of this study are that acb-tDCS produces an increase and decrease in HbO concentration in the ipsilateral and contralateral M1, respectively (Figure 2). Here, it is also worth noting that most previous studies that evaluated the effects of cb-tDCS on M1 excitability relied solely on group level responses with no control condition. However, in this study, we also investigated the inter-individual variability of acb-tDCS-induced effects on M1 excitability by establishing a sham condition.

### acb-tDCS effects at the group level

In this study, we hypothesized that acb-tDCS with the anode placed over the right cerebellum would result in decreased HbO concentration (an inhibitory effect) in the contralateral M1. At a group level, the results were consistent with our assumption, as subjects experienced an inhibitory effect in the contralateral M1 after acb-tDCS when compared to sham stimulation.

Acb-tDCS has been shown to increase Purkinje cell excitation, resulting in inhibitory output from deep cerebellar nuclei and a decrease in facilitation of thalamic cortical structures [14, 17]. In this study, we provide insight into the hemisphere-based hemodynamic effects. Ates et al. [19] in 2018 demonstrated the relationship in the excitability between the M1 and the cerebellum. By modulating cerebellar excitability using acb-tDCS and assessing changes in M1 excitability, they demonstrated that acb-tDCS has both excitatory and inhibitory effects on M1 excitability. Moreover, these results indicated the connectivity between the cerebellum and the M1 is more complex than just excitatory or inhibitory connections. Notably, similar to our study, they positioned the anode over the right cerebellum for acb-tDCS, making this a worthy comparative study. Therefore, the major contribution of our study is that we distinguished M1 excitability at the excitatory and inhibitory levels and showed that acb-tDCS has an inhibitory effect on the contralateral M1 and an excitatory effect on the ipsilateral M1 (see Figure 4(c)).

Another notable finding of this study is that ipsilateral M1 exhibited a significant increase (*p* < 0.05) in the HbO response by acb-tDCS, which was not included in our original hypothesis (see Figure 4). Similarly, Liebrand et al. have reported an increase in BOLD activity on the ipsilateral M1 by acb-tDCS with fMRI in the healthy population [20]. Interhemispheric inhibition (IHI) between the M1 cortices, a well-reported phenomenon in both healthy and stroke patients [34, 35] may be a mechanism behind this finding. Franca et al. [36] have speculated that stimulation of the cerebellum can modulate the whole brain network, including ipsilateral M1 cortex through IHI, and our findings support this hypothesis.

### Individual variability

In general, it has been shown that the responses of tDCS vary significantly between subjects and thus large standard deviations have been reported in both behavioral and imaging studies [37]. In this study, participants also demonstrated inter-individual variability for both the real and sham acb-tDCS conditions. Of the 16 participants, 7 (44%) demonstrated both expected responses simultaneously: inhibitory on the contralateral M1 and excitatory on the ipsilateral M1. Twelve (72%) demonstrated inhibitory behavior on only the contralateral M1 (Figure 5), while 11 (69%) demonstrated excitatory behavior on only the ipsilateral M1 (Figure 6). These findings are consistent with the existing literature [38], where the same qualitative results due to anodal tDCS of the M1 were reported. Their results were surprisingly variable: after anodal tDCS, about three-quarters of individuals showed the expected result (excitatory) and one quarter showed the opposite (inhibitory). These disparities are most likely due to differences in individual anatomy and neurophysiology. According to Krause et al. [39], the brain perpetuates an evenness of GABA and glutamate levels, which allows for the regulation of cortical excitability and the initiation of cortical plasticity. However, the balance of an individual’s baseline inhibition/excitation is likely to vary along a spectrum from relative inhibition to relative excitation. Therefore, the resultant balance of inhibition/excitation following tDCS would equally fluctuate between individuals. In view of this, some individuals may achieve an optimal inhibition/excitation balance with similar tDCS protocols, while others can achieve the same optimal inhibition/excitation for induction of plasticity by receiving either an Insufficient or excessive amount of stimulation.

Another explanation for this variability in results could be the fact that tDCS mostly changes the excitability in active neurons and likely has no effect on resting neuronal populations [40]. Furthermore, the excitatory or inhibitory responses totally depends upon the direction of current flow (soma to dendrites or vice versa) and axonal orientation (parallel or perpendicular) relative to the electric field [41]. Because these cerebellar neurons are not uniformly oriented, it is difficult to know which part of the neuron (axon, dendrite, or soma) is involved in these neuromodulatory effects [42]. Therefore, it is difficult to propose a particular model to identify the consistent effects of acb-tDCS.

Another interesting finding in this study was that 25% (4 out of 16 subjects) of all sham tDCS sessions had an abnormally increasing effect. An increase in excitability in the sham condition has also been reported previously [43]. However, this conspicuous increase in sham response, similar to the real tDCS response, needs to be discussed further. Previous studies investigated the sham condition with real tDCS effects using motor evoked potential (MEP) measures [44], but most did not present individual data [45-47]. Only a few studies included a sham condition and presented individual responses as well [43, 48]. Horvath et al. (2016) presented noticeably higher variability for the sham condition at the individual level. They demonstrated that 38% of the sham sessions exhibited an abnormal increase in MEP amplitude when compared to the baseline values [43]. Another study found that 28% of all sham sessions showed an increasing or decreasing effect of MEP amplitude [49]. Dyke et al. (2016) observed differences for sham stimulation between the TMS recruitment curve slope and the baseline in some of their individuals [48]. Our study reports that hemodynamic responsiveness can be induced with sham tDCS in a few healthy subjects in accordance with previous studies, although the underlying mechanism of responsiveness to sham stimulation remains unclear. Further study is needed to elucidate the individual-level variability of sham tDCS.

### Limitations and future directions

In this section, we would like to draw attention to some limitations. First, although we advised the subjects to stay awake throughout the experiment, we were unable to standardize their wakefulness or drowsiness levels. Therefore, some subjects may have had varying levels of drowsiness and wakefulness that resulted in individual disparities in their brain states. Second, the cortical excitability of the M1 was not measured in this study, thus we cannot confirm whether acb-tDCS could induce changes in corticospinal tract excitability. However, as Takano et al. have previously demonstrated that acb-tDCS elicits a decrease in corticospinal excitability after 13 min of stimulation [50], we could infer that our protocol (20 min of acb-tDCS) was sufficient to induce changes in corticospinal excitability. Third, our experiment did not include behavioral measurements, hindering us from investigating whether the changes in HbO on the M1 cortices are related to behavioral outcomes. Future research, including cortical excitability measurement and behavioral scales, is needed to develop rigorous evidence of CBI as well as its effect on behavioral function.

## Conclusion

The acb-tDCS induces hemodynamic changes in the M1 (contralateral side: decrease in HbO, ipsilateral side: increase in HbO), which could be explained by CBI and interhemispheric inhibition. However, individual variabilities of hemodynamic changes in the M1 in response to acb-tDCS were found, and further study is needed to investigate whether these variabilities affect the clinical effect of cb-tDCS over multiple sessions and in patients with neurological diseases such as stroke and Parkinson’s disease.

## Abbreviations

CBI: cerebellar brain inhibition
M1: primary motor cortex
fNIRS: functional near-infrared spectroscopy
acb-t DCS: anodal cerebellar transcranial direct current stimulation
HbO: oxygenated hemoglobin
tDCS: transcranial direct current stimulation
cb-tDCS: cerebellar direct current stimulation
fMRI: functional magnetic resonance imaging
SPECT: single-photon emission computerized tomography
PET: positron emission tomography
HBR: deoxygenated hemoglobin

## Acknowledgment

None

